# Dominant Formation of DNA:RNA Hybrid G-Quadruplexes in the Genomes of Living Human and Mouse Cells

**DOI:** 10.1101/2024.12.27.630557

**Authors:** Zheng Tan

## Abstract

Guanine-rich nucleic acids can form four-stranded structures known as G-quadruplexes (G4s), which play critical roles in essential cellular activities. Putative G-quadruplex-forming sequences (PQSs) are abundant in the genomes of animal cells. To date, research on G4s has focused primarily on intramolecular G4s formed by four contiguous G-tracts from the same nucleic acid strand. However, our recent study indicates that a hybrid type of G4s, DNA:RNA hybrid G4s (hG4s), comprising G-tracts from both DNA and RNA, is, in fact, the most abundant G4 species found in the genomes of living yeast cells. Here, we demonstrate that hG4s also readily form and predominate in the genomes of living human and mouse cells, with an abundance several orders of magnitude greater than that of intramolecular DNA G4s (dG4s). Furthermore, each individual G-tract in a PQS tends to form an hG4 independent of adjacent G-tracts. This independence of DNA G-tracts is maintained even in PQSs containing four G-tracts, despite their potential to form intramolecular DNA G4s only by themselves. Collectively, our studies demonstrate that hG4s are the predominant physiological G4 structures in the genomes of eukaryotic organisms. Given that hG4s differ significantly from intramolecular dG4s in terms of structural form, formation mechanism, abundance, and association with transcription, there is a clear need for a comprehensive revision of our current understanding of genomic G4s and their functional roles. We hypothesize that hG4 formation may serve as a global sensing mechanism to monitor RNA levels in cells to maintain transcriptional homeostasis.

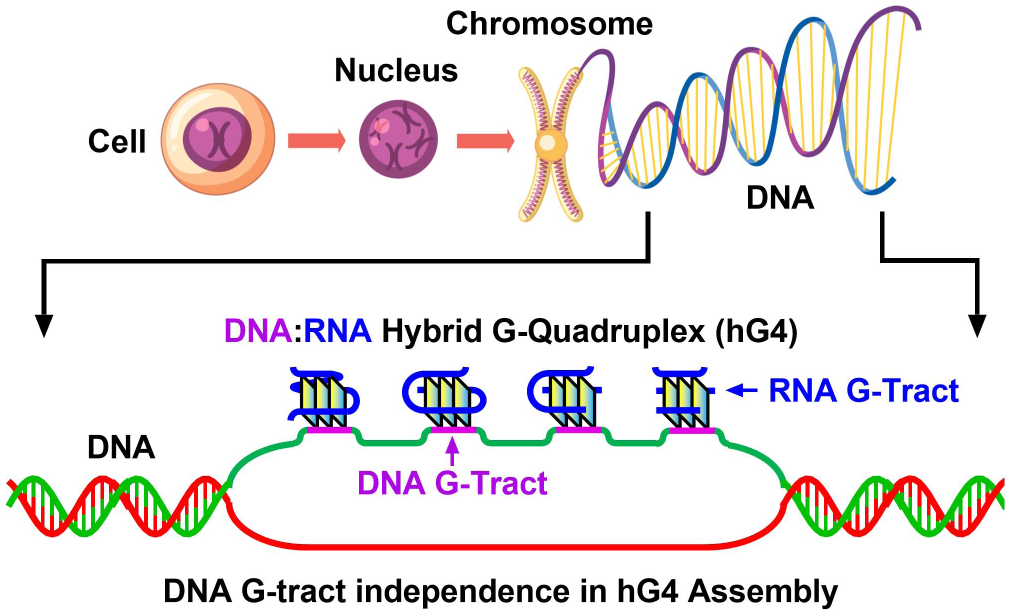

## Introduction

G-quadruplexes (G4s) are four-stranded, non-canonical secondary structures formed by guanine-rich nucleic acids. The most commonly studied G4s consist of three or more stacked G-tetrads, each comprising four guanine residues connected in a square planar configuration by Hoogsteen hydrogen bonds. G4s have been implicated in various biological processes, including transcription, replication, genome stability, and epigenetic regulation. Putative G-quadruplex-forming sequences (PQSs) are particularly enriched in promoter regions, suggesting a role in the regulation of gene expression (1). For instance, G-quadruplex formation in the promoter region of the c-myc gene is known to regulate its expression (2), making it a potential target for cancer therapy.

PQSs are abundant in the genomes of eukaryotic cells (3). The canonical PQSs, which are capable of forming intramolecular DNA G4s (dG4s), can be described by the following sequence consensus: G_≥3_N_1-7_G_≥3_N_1-7_G_≥3_N_1-7_G_≥3_, where N denotes any nucleotide, and the numbers indicate the range of nucleotides that may be present between the G-tracts (4). In the human genome, approximately 370,000 such PQSs have been identified based on this consensus (4). Furthermore, experimental data demonstrate that G4s can form in PQS motifs with longer loops (5) or with fewer than three guanines per G-tract (6), as well as in sequences that do not strictly conform to the PQS consensus (7,8). Computational and deep sequencing approaches have identified over 700,000 regions in the human genome that could potentially fold into G4 structures (9). The G4s mentioned above, which have been the focus of G4 biology, are intramolecular G4s with four continuous G-tracts from the same strand. Their structural forms have mainly been studied under simplified in vitro conditions.

However, G4 formation in the genomes of living cells occurs in a distinctly different environment that influences various aspects of G4s, including formation kinetics, conformation, stability, and interactions with other molecules (10). Consequently, the structural forms of G4s in the genomes of living cells have remained largely unclear. Our previous studies have shown that hybrid G4s (hG4s), composed of G-tracts from both DNA and RNA, can form in both linear double-stranded DNA and plasmids transcribed in vitro (6,11–15), as well as in plasmids transfected into bacterial cells (16). Recently, we have rigorously demonstrated that such hG4s also readily form as the predominant G4 species in the genomes of living yeast cells (17) using an Okazaki fragment (OKF) extension stop assay. Since yeast is a unicellular organism, it remains unknown whether hG4s would form in much more complex multicellular organisms. To address this concern, we examined the structural forms of G4s in the genomes of living human and mouse cells. Our results indicate that hG4s are the predominant G4 species, primarily assembled at single DNA G-tracts.

## Materials and Methods

### Identification of PQS and control motifs in the genome

Genome sequence files in fasta format and chromosome size files for human (hg38) and mouse (mm10) were downloaded from the UCSC Genome Browser (http://genome.ucsc.edu/) using the URLs provided on the website. PQSs defined by the consensus G_≥2_(N_1-7_G_≥2_)_≥0_ and G_≥3_(N_1-7_G_≥3_)_≥3_ were identified using an regular expression G{2,}(.{1,7}?G{2,}){0,} and G{3,}(.{1,7}?G{3,}){3,}, respectively, as described, respectively (17), with the output saved in bed files. Control motifs were generated by shuffling the corresponding PQSs to random genomic locations.

### Identification of orphan PQSs

Orphan PQSs with a G_≥2_-free region on either side in each strand were identified using the original PQS bed files prior to strand splitting as described (17).

### Sequencing data from the Gene Expression Omnibus (GEO) database

The original paired-end sequencing data of the OKFs, G4P-ChIP, and G4access were downloaded in fastq format from NCBI’s GEO database (http://www.ncbi.nlm.nih.gov/geo/) using the fasterq-dump command with the --split-3 option, along with the accession numbers listed in Table 1.

**Table 1.** Sequencing data downloaded from GEO.

| Organism | G4 Probe | GEO Accession | Cell Line | Reference |
| --- | --- | --- | --- | --- |
| Human | OKF | GSE239856 | FT194 | (18) |
|  | G4P | GSE133379 | A549 | (19) |
|  | G4access | GSE187007 | HaCaT, K562, Raji | (20) |
| Mouse | OKF | GSE142996 | mESC E14 | (21) |
|  | G4P | GSE133379 | 3T3 | (19) |
|  | G4access | GSE187007 | Embryonic stem cell | (20) |

### Distribution of OKFs, G4P, and G4access signals across PQSs

The downloaded fastq files were processed using the fastp tool (22) to trim adapters, remove duplicate reads, and filter out reads of low quality or shorter than 10 nucleotides (nts) using default settings. The resulting fastq files were then aligned to the human (hg38) or mouse (mm10) genome using Bowtie2 with the following options: --sensitive-local --no-unal --no-mixed --no-discordant. The aligned reads were then filtered using samtools view with the options: -h -f 3 -q 20 -F 0x100 and the results were saved in bam format. For G4P and G4access data, bigwig files were generated from the bam files as described (17,19). For OKFs, the resulting OKF bam files were converted to bed files using a custom script and then to bedgraph and finally to bigwig files for the 3’ ends as described (17). Profiling of OKF 3’ ends, G4P, and G4access signals across PQSs was performed with the bigwig files as described previously (17). If a PQS bed file was too large, 200,000 PQSs were randomly sampled for profiling to reduce computational time. The final profiling result was the average of all samples in each dataset.

## Results

### Abundance of PQSs capable of forming hG4s

In principle, a PQS in a genome can form either an intramolecular dG4 or multiple hG4s with different DNA:RNA G-tract ratios depending on the number of G-tracts it possesses (Figure 1A). Based on our recent study (17), hG4 formation in a genome can occur in PQSs containing one or more G-tracts, each consisting of two or more contiguous guanines. Following this rule, we first used the consensus G_≥2_(N_1-7_G_≥2_)_≥0_ to identify such PQSs in the human and mouse genomes. In addition, canonical intramolecular PQSs with four G-tracts of three or more guanines were identified using the consensus G_≥3_(N_1-7_G_≥3_)_≥3_. As shown in Table 2, the human genome has 152,656,667 G4-forming PQSs, including hG4-forming PQSs, which is 391 times the number of canonical PQSs that can form intramolecular dG4s. The mouse genome has 135,593,514 G4-forming PQSs when the hG4-forming PQSs are included, which is 275 times the number of canonical PQSs that can form intramolecular dG4s.

**Figure 1.**
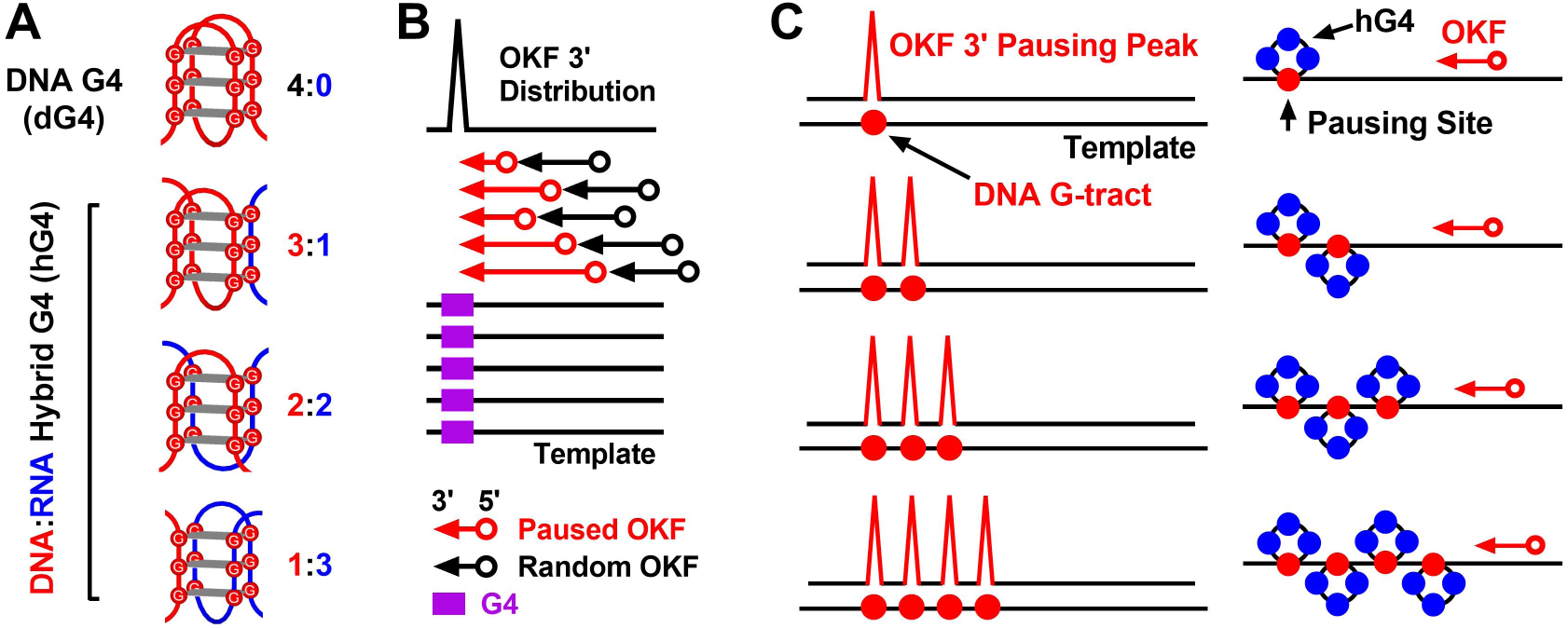
Representative examples of hG4 assembly in the genome and detection by the OKF extension stop. (A) G-tract assembly of a canonical intramolecular DNA G4 (dG4) and hybrid DNA:RNA G4s (hG4s). (B) Alignment of the 3’ ends of the paused OKF extensions to G4s, resulting in pausing peaks at the PQSs. (C) The number of pausing peaks indicates hG4 assembly in each individual DNA G-tract (right).

**Table 2.** Number of classified PQSs in the human and mouse genomes and the form of G4s they can form.

| G4 Type | Name* | G-tetrads | Consensus | Count (Human) | Count (Mouse) |
| --- | --- | --- | --- | --- | --- |
| dG4 | 4G3+ | $\geq 3$ | $G_{\geq 3}(N_{1-7}G_{\geq 3})_{\geq 3}$ | 389,584 | 491,854 |
| hG4/dG4 | 4G2+ | $\geq 2$ | $G_{\geq 2}(N_{1-7}G_{\geq 2})_{\geq 3}$ | 9,130,747 | 6,535,901 |
| hG4 | 3G2+ | $\geq 2$ | $G_{\geq 2}(N_{1-7}G_{\geq 2})_2$ | 10,598,355 | 8,813,893 |
| hG4 | 2G2+ | $\geq 2$ | $G_{\geq 2}N_{1-7}G_{\geq 2}$ | 30,110,037 | 27,667,418 |
| hG4 | 1G2+ | $\geq 2$ | $G_{\geq 2}$ | 102,817,528 | 92,576,302 |
| hG4/dG4 | 1234G2+ | $\geq 2$ | $G_{\geq 2}(N_{1-7}G_{\geq 2})_{\geq 0}$ | 152,656,667 | 135,593,514 |

\*The digit prefixing the “G” denotes the number of G-tracts, the digit plus “+” suffixing the “G” denotes the minimal size of the G-tracts. 4G3+ represents canonical intramolecular G4s with three or more G-tetrads.

### Detection of hG4 formation by OKF extension stop assay

G4s act as roadblocks (23), physically impeding protein translocation along a DNA strand (24). Our recent study demonstrated that OKF synthesis in yeast pauses at both hG4 and dG4, and the extension of OKF 3’ ends reports G4 formation at PQSs in the genomes with pausing peaks in their profile (17) (Figure 1B). A notable finding of that study was that the number of major pausing peaks consistently matched the number of G-tracts in the PQSs (Figure 1C, left side). These facts suggest that a DNA:RNA G-tract ratio of 1:3 is the preferred configuration for hG4 assembly in all cases, regardless of the number of G-tracts in a PQS (Figure 1C, right side).

To investigate whether hG4s could form in the human and mouse genomes as in the yeast, we first downloaded human (18) and mouse (21) OKF sequencing data from the GEO database (Table 1) and profiled the OKF 3’ ends around three groups of orphan PQSs, each containing a single stretch of GG, GGG, or GGGG flanked on either side by a G_≥2_-free region of at least 20 to 100 nts (Figure 2). The profiles obtained all showed a prominent single peak at the PQSs (black arrowhead) on the PQS-bearing template strand, indicating the formation of hG4s at the PQSs with one G-tract from DNA and three from RNA. Notably, the peak amplitude remained largely unchanged relative to the background as the G-tract-free region increased from 20 to 100 nts (panels from left to right). This feature excluded the possibility that the pause was caused by the orphan PQSs recruiting neighboring G-tracts to form intramolecular dG4s rather than hG4s. Conversely, the peak amplitude increased as the size of the G-tracts increased from 2 to 4 nts (panels from top to bottom), which is consistent with the known fact that G4s with longer G-tracts are more stable (25).

**Figure 2.**
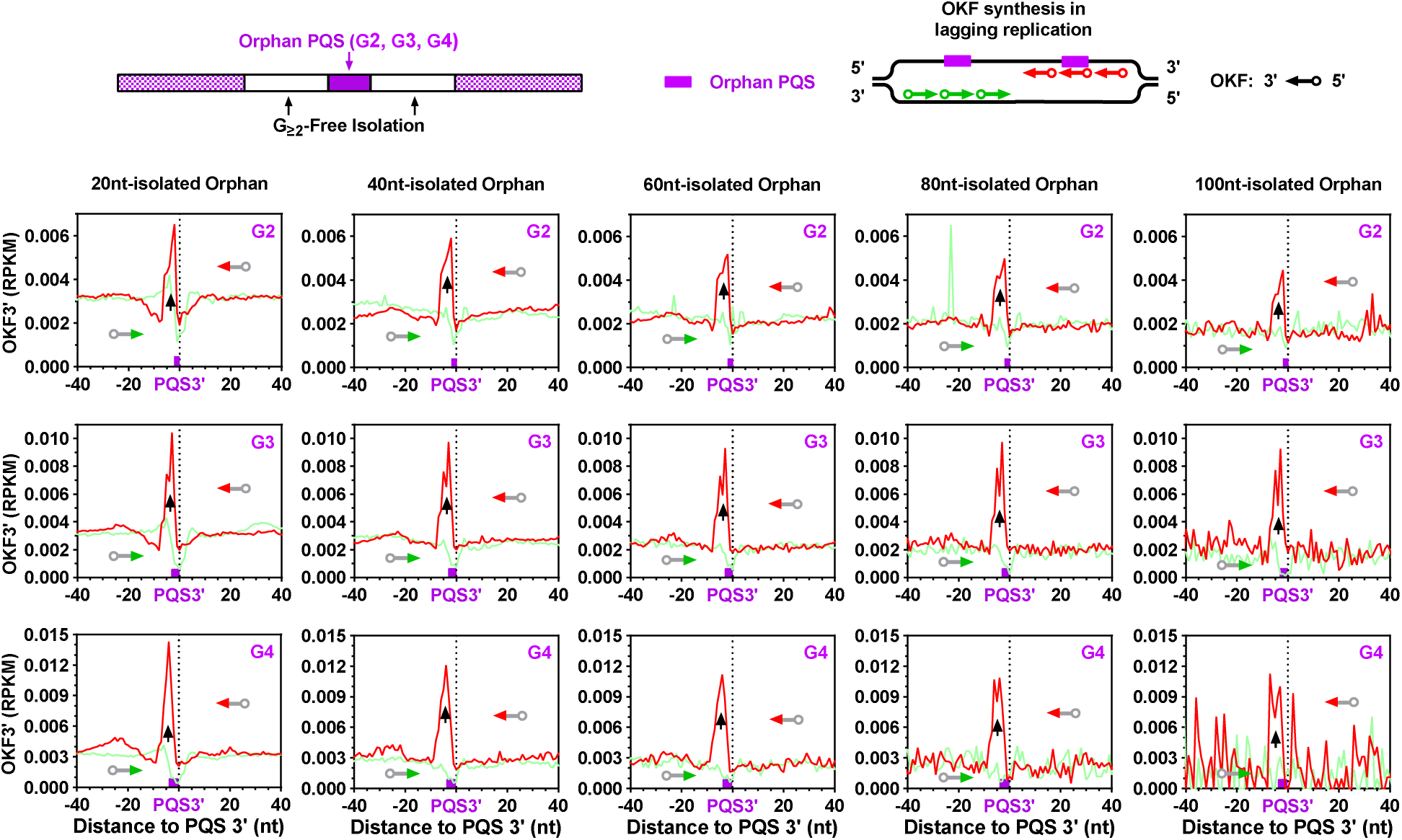
Detection of hG4 formation in the human genome by OKF 3’ end pausing at orphan PQSs consisting of a single GG, GGG, or GGGG tract isolated from adjacent G-tracts by at least the specified number of nucleotides (nts). Bin size: 1 nt.

We next examined the 3’ end pausing of OKFs in three groups of regular PQSs, each containing two G-tracts of the same size and loops ranging from 1 to 7 nts (Figure 3). Two pausing peaks were observed, similar to those in yeast. Importantly, the distance between the two peaks increased as the size of the loops between the G-tracts increased from 1 to 7 nts (panels from top to bottom), indicating that the pausing was indeed caused by the formation of hG4 at each G-tract. Theoretically, these PQSs could form a single hG4 by recruiting two RNA G-tracts, resulting in a single OKF 3’ pausing peak. The lack of such behavior suggests that each G-tract instead intended to recruit RNA G-tracts independently.

**Figure 3.**
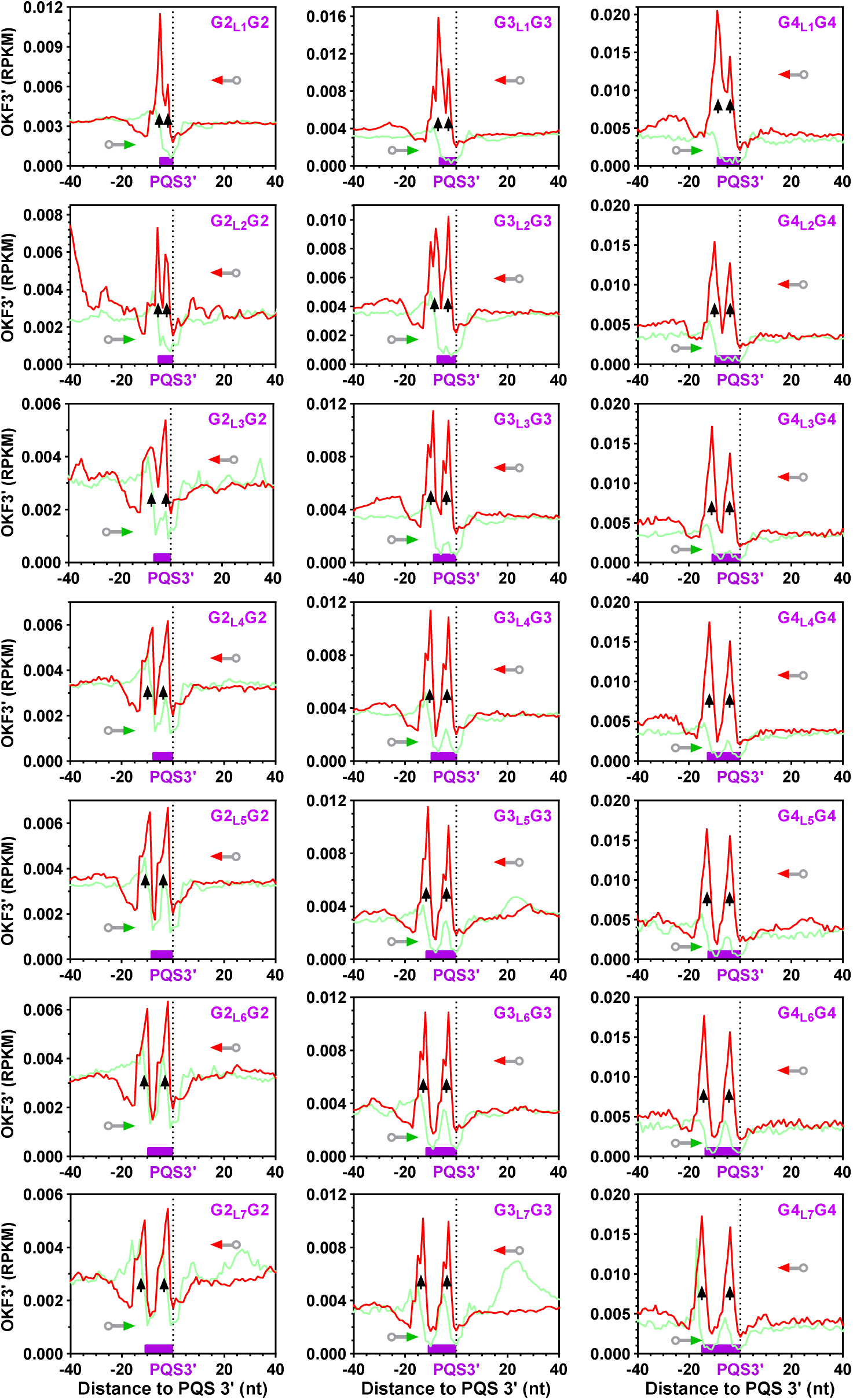
Detection of hG4 formation in the human genome by OKF 3’ end pausing at PQSs containing two G-tracts of different sizes along with loops of increasing length. The sizes of the G-tracts and loops are given in the PQS sequences in the panels, with numbers appended to “G” and “L”, respectively. Bin size: 1 nt.

We also examined three groups of regular PQSs, each with three G-tracts of the same size and loops ranging from 1 to 7 nts (Figure 4). Accordingly, three pausing peaks were observed, and the distances between them also increased with increasing loop size (panels from top to bottom). When PQSs with four G_2_ tracts were examined, the number of pausing peaks also matched the number of G-tracts, and the distance between the peaks increased with increasing loop size (Figure 5). In this particular scenario, the PQSs are capable of forming intramolecular dG4s on their own. However, the well-isolated peaks strongly suggest that hG4s with a DNA:RNA G-tract ratio of 1:3 were preferentially formed rather than the intramolecular dG4s. Otherwise, a single pausing peak would be expected. Taken together, the equality between the number of peaks and the number of G-tracts indicated a general and dominant independence of the G-tract in the hG4 assembly.

**Figure 4.**
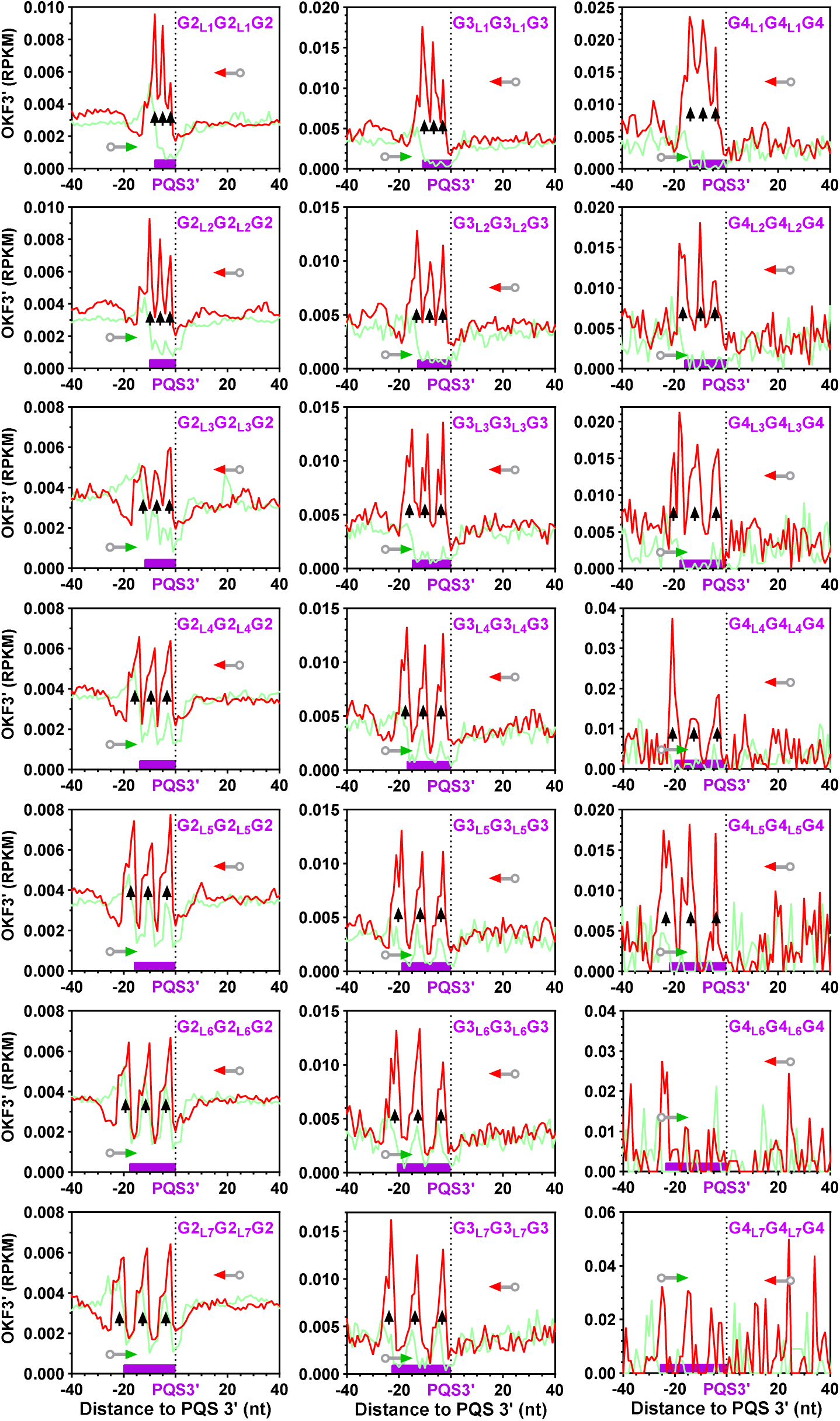
Detection of hG4 formation in the human genome by OKF 3’ end pausing at PQSs containing three G-tracts of different sizes along with loops of increasing length. The peaks became less obvious or disappeared when the number of PQSs became too few due to deterioration of the signal-to-noise ratio. Bin size: 1 nt.

**Figure 5.**
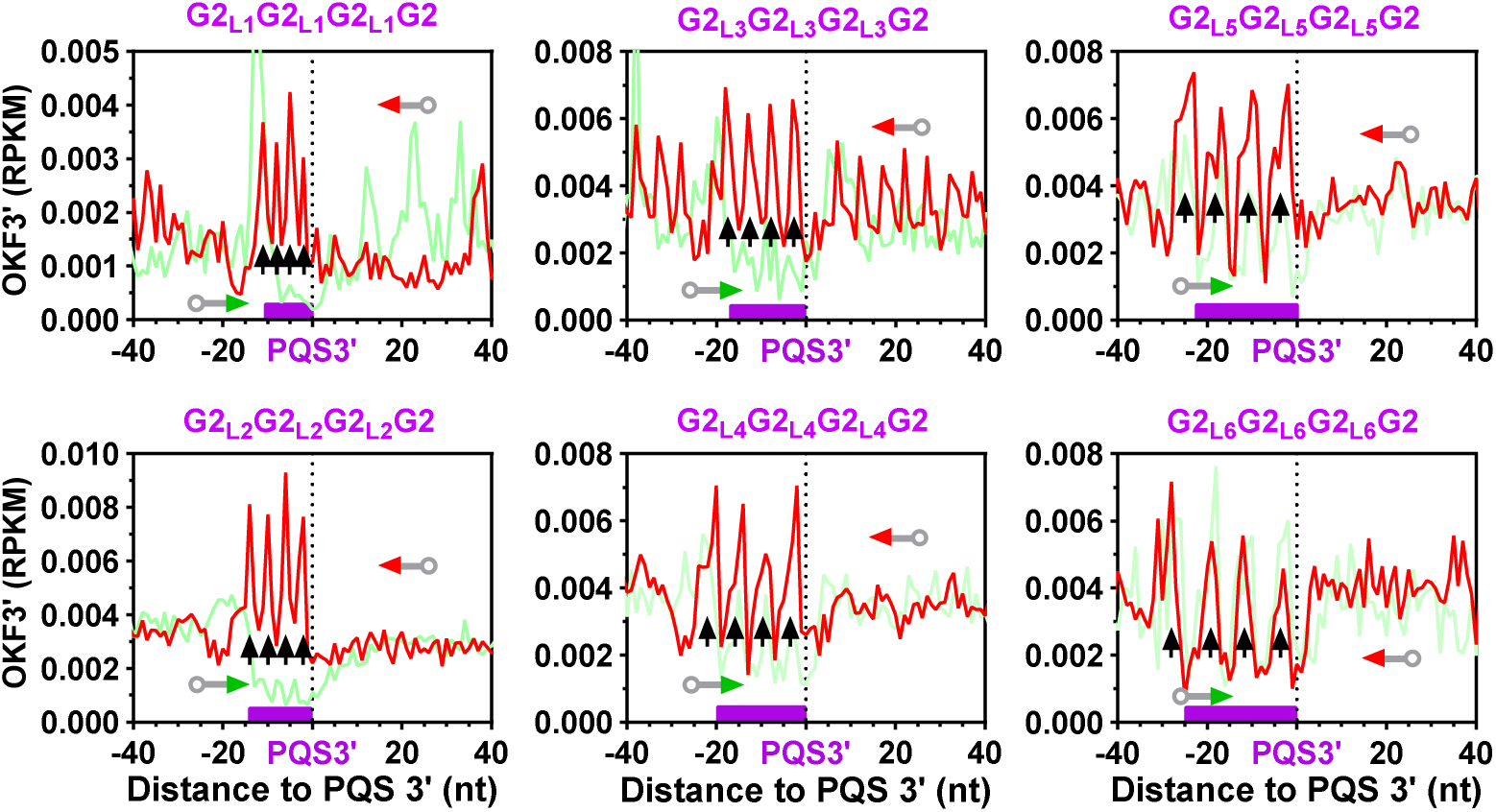
Detection of hG4 formation in the human genome by OKF 3’ end pausing at PQSs containing four G-tracts of different sizes along with loops of increasing length. Bin size: 1 nt.

### Detection of hG4 formation with G4 probe (G4P) protein

We have previously developed an artificially engineered G4 probing (G4P) protein with high affinity and specificity for G4s (Figure 6A). This protein recognizes both hG4 and dG4 with two G4 recognition domains and was expressed in human, mouse cell lines (19), and yeast (17) to detect G4 formation in the genomes using the ChIP-seq technique. To test whether G4s can form at PQSs with fewer than four DNA G-tracts in the human genome, we profiled the G4P signal over the PQSs capable of forming either dG4s or hG4s (Figure 6). As expected, the 4G2+ PQSs containing four consecutive G-tracts of two or more guanines and capable of forming intramolecular dG4s were bound by G4P, as indicated by the enrichment of G4P at the PQSs (panel B). In addition, those with only one to three G-tracts (panels C-E) were also bound by G4P. Since these PQSs are not able to form dG4s by themselves, the results support the formation of hG4s in these PQSs. The enrichment disappeared when the PQSs were shuffled to random locations in the genome, indicating a dependence on the PQSs.

**Figure 6.**
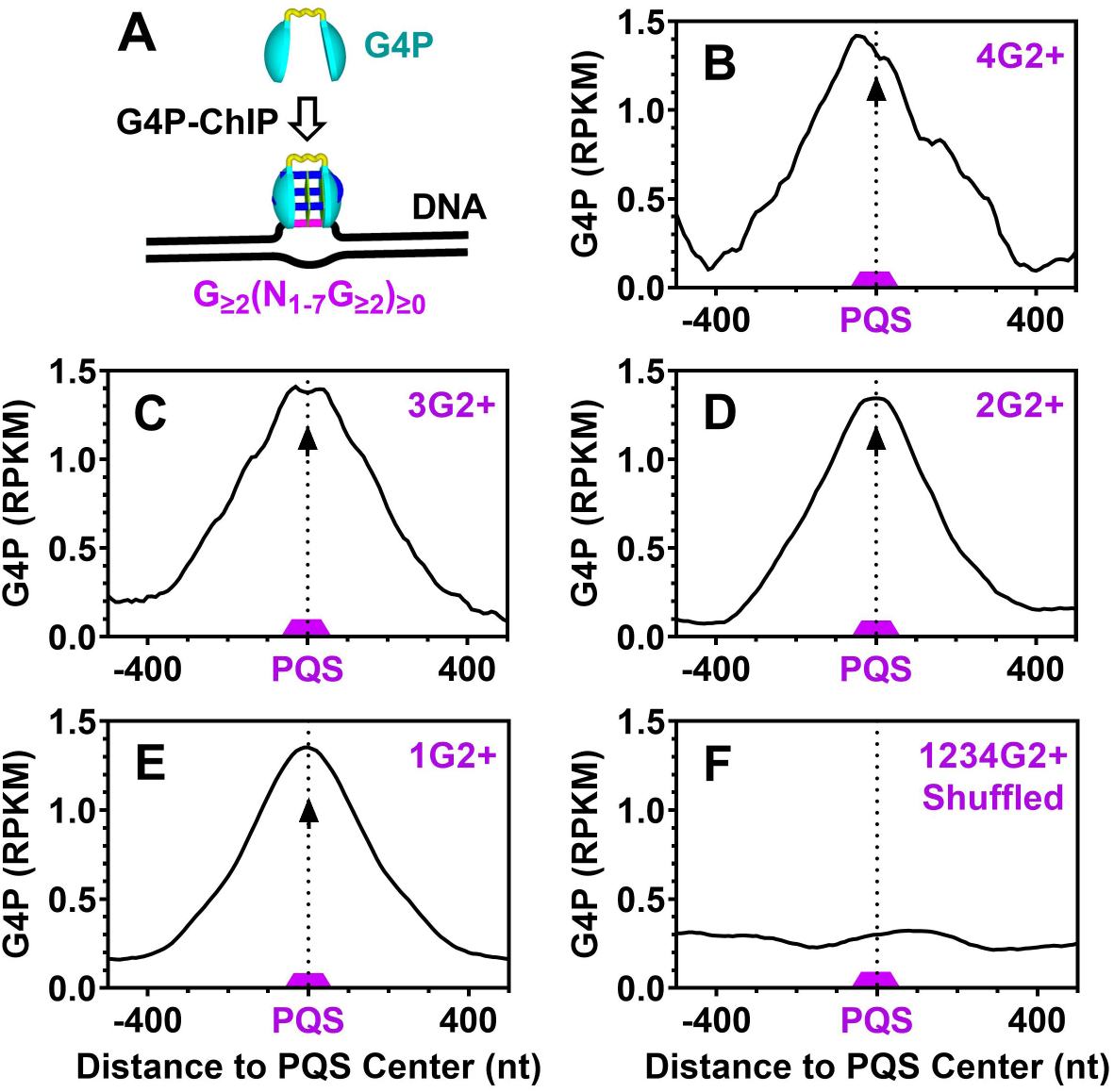
Detection of hG4 formation in the human genome by G4P at orphan PQSs containing 1 to 4 or more G_≥2_ tracts that can form G4s of two or more G-tetrads. (A) Schematic illustration of G4 detection by G4P ChIP-Seq. (B-E) Enrichment of G4P at PQSs capable of forming (B) dG4s or (C-E) hG4s. (F) Distribution of G4P at randomly shuffled PQSs. The PQSs were flanked on both sides by at least 50 nt of G_≥2_-free regions. Bin size: 10 nt.

### Detection of hG4 formation with G4access

To further confirm the formation of G4s at the PQSs, we used sequencing data obtained using the G4access technique (20). This technique is designed to detect G4s based on their resistance to micrococcal nuclease (MNase) hydrolysis in open chromatin (Figure 7A) and provides significantly better resolution compared to ChIP-seq. In support of the presence of G4s at the PQSs, a G4access peak was observed at these PQSs (Figure 7, B-E). Notably, the amplitude of the peak increased with the number of G-tracts in the PQSs, indicating protection of more G-tracts within the G4s. Furthermore, the PQS dependence of G4 formation was confirmed by the disappearance of the G4access signal when the PQSs were shuffled to random locations (Figure 7F).

**Figure 7.**
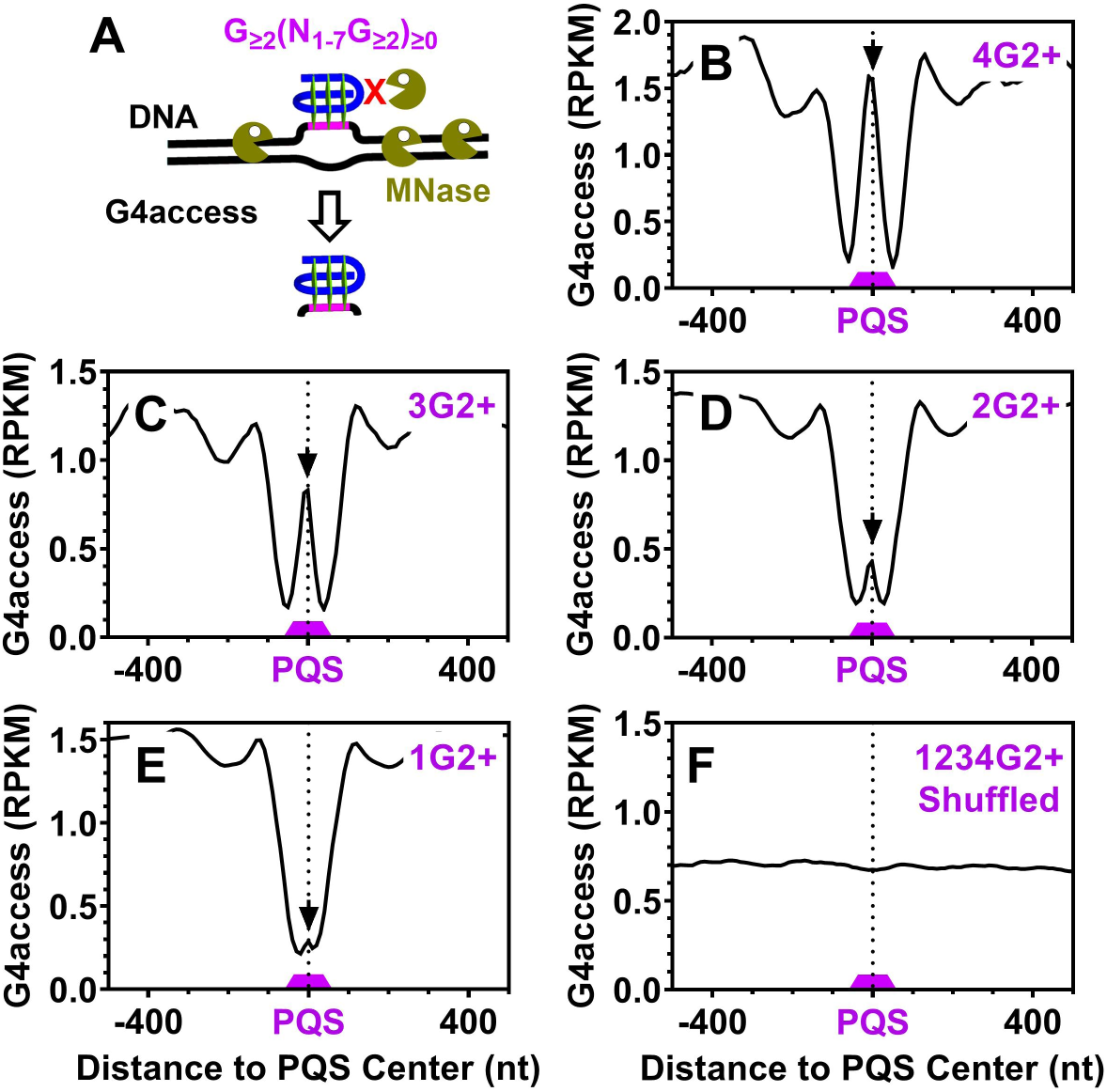
Detection of hG4 formation in the human genome by G4access at orphan PQSs containing 1 to 4 or more G_≥2_ tracts that can form G4s of two or more G-tetrads. (A) Schematic illustration of G4 detection using G4access. (B-E) Enrichment of G4access signal at PQSs capable of forming (B) dG4s or (C-E) hG4s. (F) Distribution of G4access signal at randomly shuffled PQSs. The PQSs were flanked on both sides by at least 50 nt of G_≥2_-free regions. Bin size: 10 nt.

The results shown in Figure 2 to Figure 7 are for human cells. The same set of analyses was also performed in exactly the same way for mouse cells using data from the GEO database (19–21), and the results are shown in Supplementary Figure S1 to Figure S6. These results indicate that hG4 formation in the mouse genome behaved in the same way as in the human genome.

## Discussion

The OKF extension stop assay (17) detects G4 formation in living cells in a completely natural cellular state. Using this method, our current study of hG4 formation in the genomes of living human and mouse cells reproduced the key features we recently reported in the genomes of living yeast cells (17). First, hG4s can be efficiently formed with a single DNA G-tract of two or more guanines. Second, hG4s induce pausing of OKF extension, and the number of pausing peaks is equal to the number of G-tracts in the PQSs, indicating hG4 assembly at each individual DNA G-tract. Third, although PQSs with four G-tracts are capable of forming intramolecular dG4s by themselves, they still prefer to form hG4s at each individual G-tract. Taken together, our studies in human, mouse and yeast cells indicate that hG4s assembled by such a strategy are the overwhelmingly dominant G4 species in the genomes of eukaryotic cells.

In both our previous (17) and current studies, our results consistently showed that the peaks of the OKF 3’ ends are well resolved and that their number is equal to the number of G-tracts in the PQSs, despite the number of G-tracts in the PQSs. This feature indicates that the formation of hG4s at a DNA:RNA ratio of 1:3 occurred preferentially at each DNA G-tract, regardless of the number of G-tracts in the PQSs (Figure 1C). The formation of an intramolecular G4 in a single strand of nucleic acid may be advantageous because the loops hold the G-tracts together and the single-stranded form is more flexible, allowing it to fold back and forth easily to assemble an intramolecular G4. However, this process can be challenging for a PQS confined within a rigid DNA double helix. Incorporation of G-tracts from RNA, which is abundant in the cellular environment, appears to be a viable strategy to mitigate this challenge.

hG4s differ from canonical intramolecular dG4s in several aspects, including their structural forms, formation mechanisms (12), abundance (Table 2), kinetics, stability (26), and, most importantly, their interaction with transcription. For these reasons, hG4s are likely to have unique functional implications compared to canonical intramolecular dG4s. For instance, loop size is a critical parameter that influences intramolecular G4 formation in terms of kinetics, stability, and folding topology (5,27–31). However, in the case of genomic hG4s, loops may primarily serve as spacers to separate adjacent hG4s due to the independence of the DNA G-tract in hG4 assembly. In our view, the most important aspect is the extensive DNA-RNA interactions mediated by hG4s, which can occur at over one hundred million genomic sites in human and mouse cells. This ability allows genomic DNA to sense RNA levels and form hG4s in response, creating a feedback to regulate transcription. We propose that such a feedback loop may play a role in maintaining global transcriptional homeostasis in a cell.

## Acknowledgments

This work was supported by the National Natural Science Foundation of China (grant # 22377009 and 22037004) and the Shanxi “1331 Project”.

## Supplementary Information for

This appendix includes supplementary Figures S1-S6, analyses of hG4 formation in the mouse genome.

**Figure S1.**
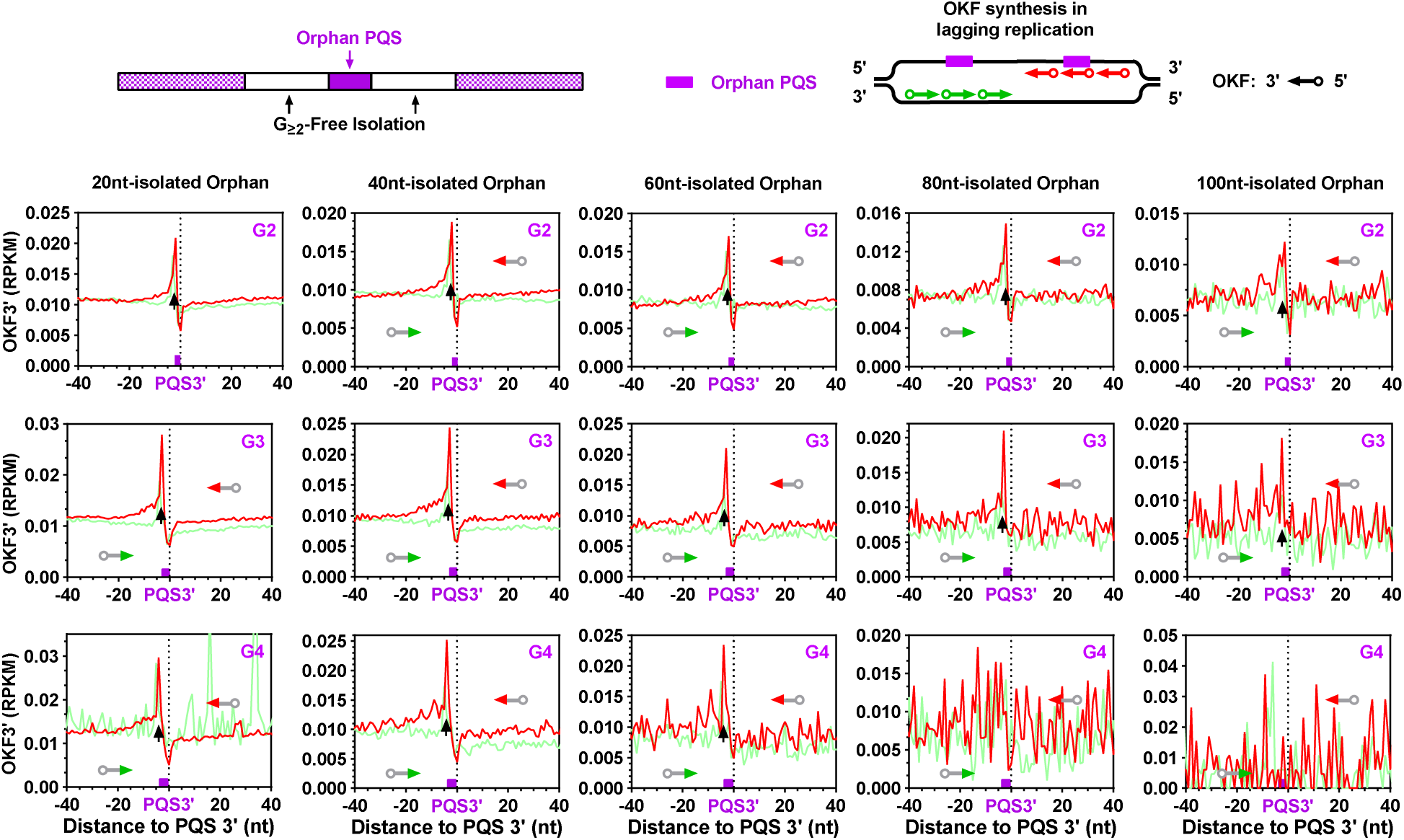
Detection of hG4 formation in the mouse genome by OKF 3’ end pausing at orphan PQSs of a single GG, GGG, GGGG tract isolated from adjacent G-tracts by at least the specified number of nucleotides (nts). Bin size: 1 nt.

**Figure S2.**
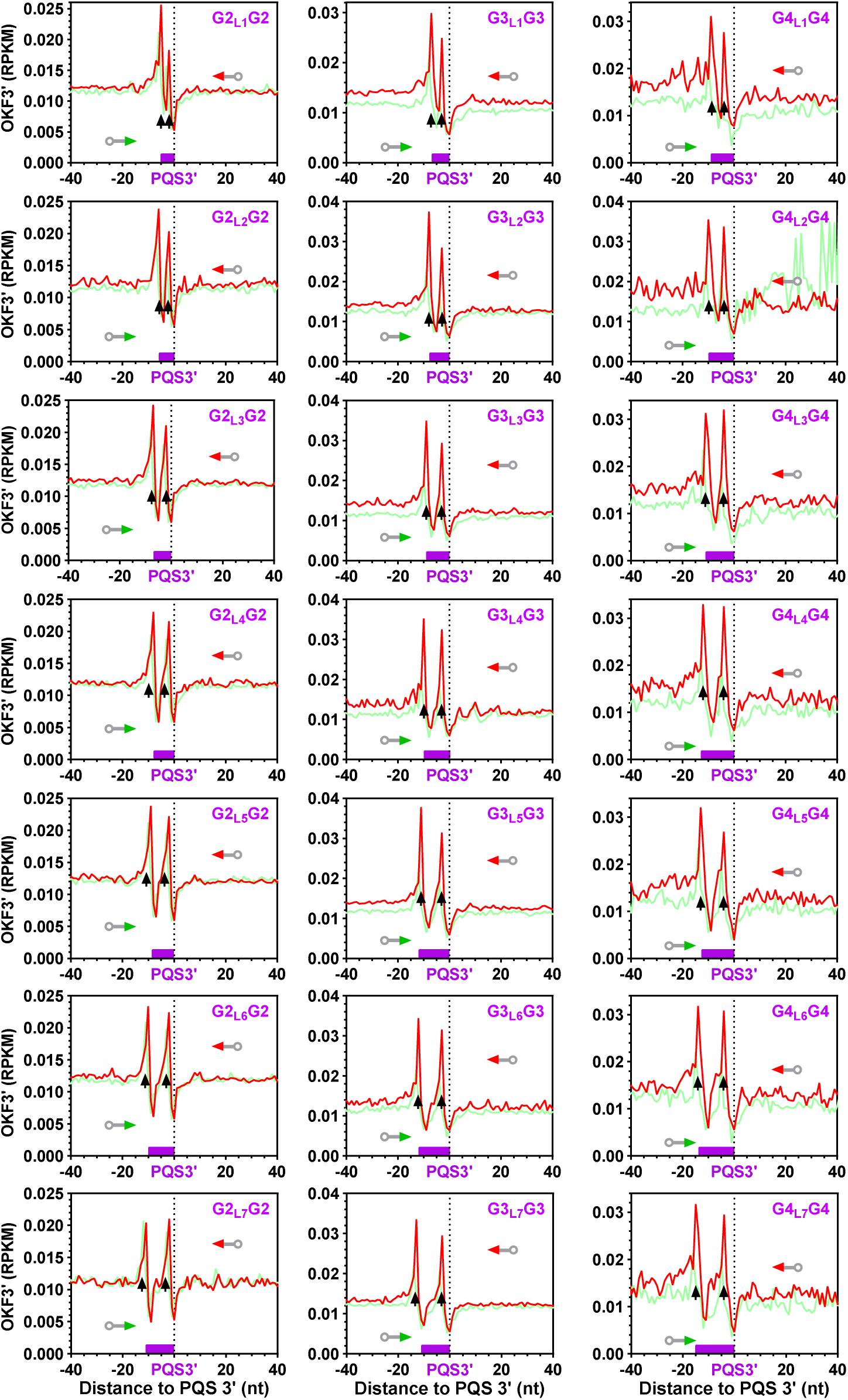
Detection of hG4 formation in the mouse genome by OKF 3’ end pausing at PQSs containing two G-tracts of different sizes along with loops of increasing length. The sizes of the G-tracts and loops are given in the PQS sequences in the panels, with numbers appended to “G” and “L”, respectively. Bin size: 1 nt.

**Figure S3.**
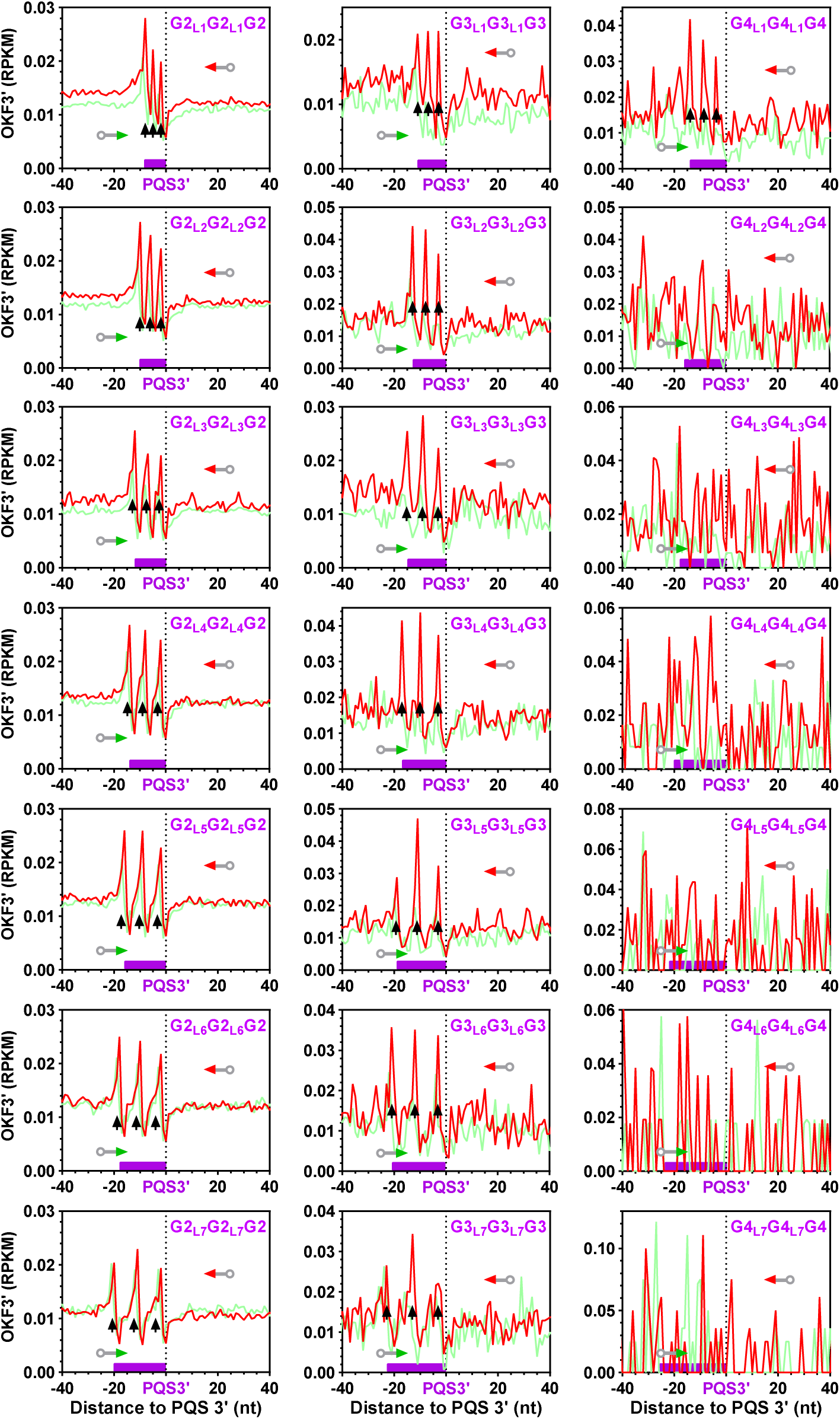
Detection of hG4 formation in the mouse genome by OKF 3’ end pausing at PQSs containing three G-tracts of different sizes along with loops of increasing length. The peaks became less obvious or disappeared when the number of PQSs became too few due to deterioration of the signal-to-noise ratio. Bin size: 1 nt.Bin size: 1 nt.

**Figure S4.**
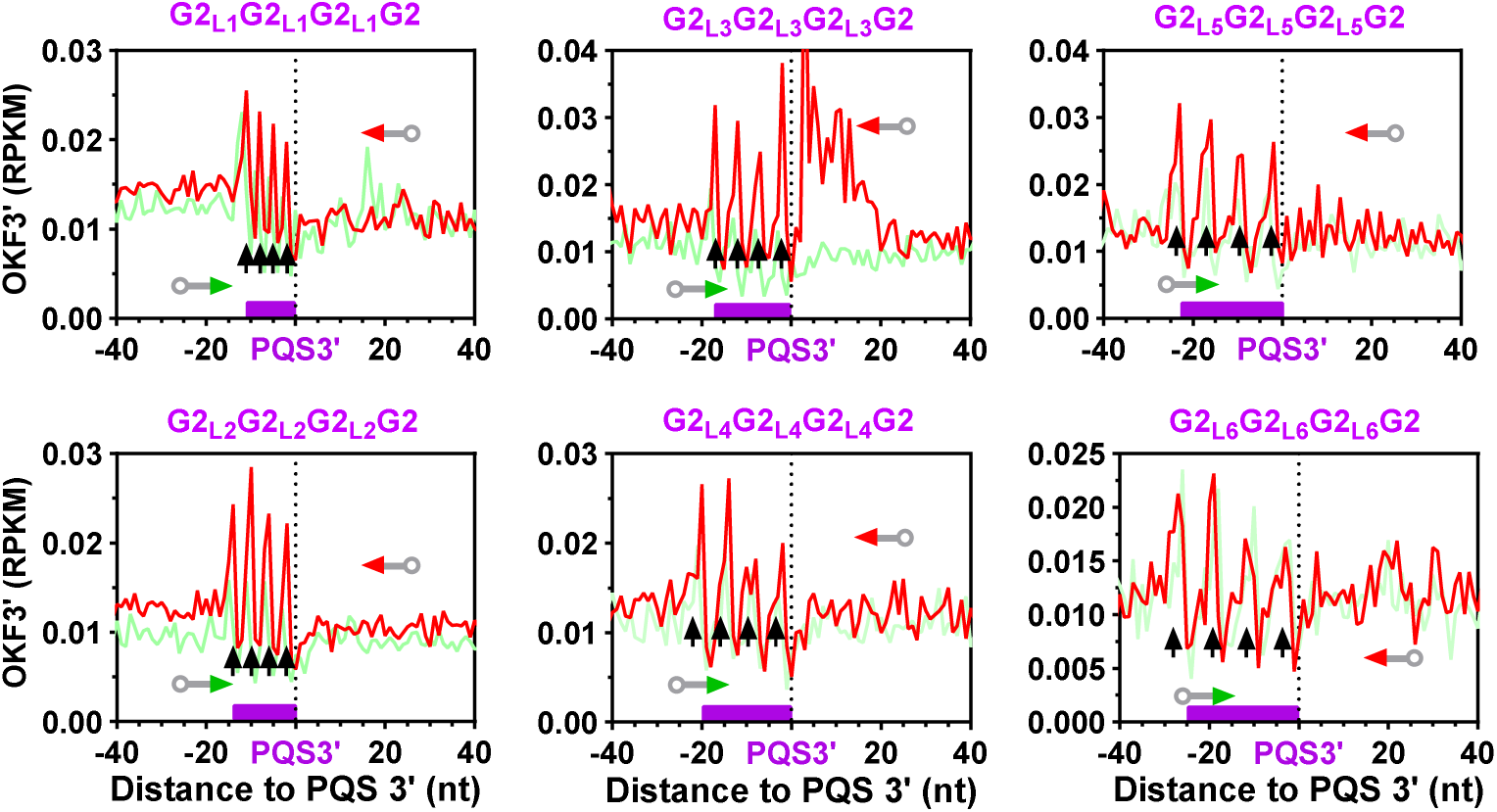
Detection of hG4 formation in the mouse genome by OKF 3’ end pausing at PQSs containing four G-tracts of different sizes along with loops of increasing length. Bin size: 1 nt.

**Figure S5.**
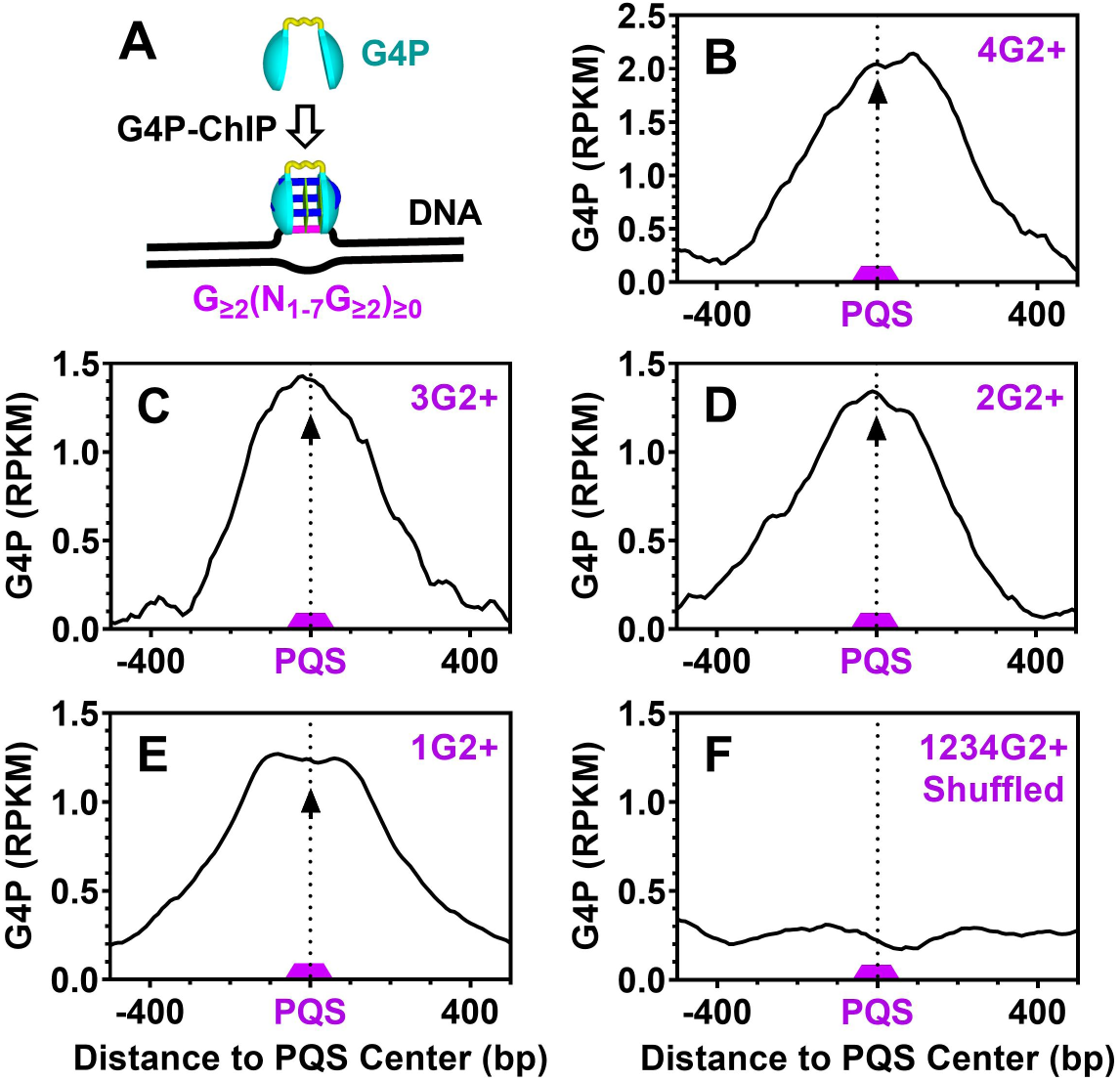
Detection of hG4 formation in the mouse genome by G4P at orphan PQSs containing 1 to 4 or more G_≥2_ tracts that can form G4s of two or more G-tetrads. (A) Schematic illustration of G4 detection by G4P ChIP-Seq. (B-E) Enrichment of G4P at PQSs capable of forming (B) dG4s or (C-E) hG4s. (F) Distribution of G4P at randomly shuffled PQSs. The PQSs were flanked on both sides by at least 50 nt of G_≥2_-free regions. Bin size: 10 nt.

**Figure S6.**
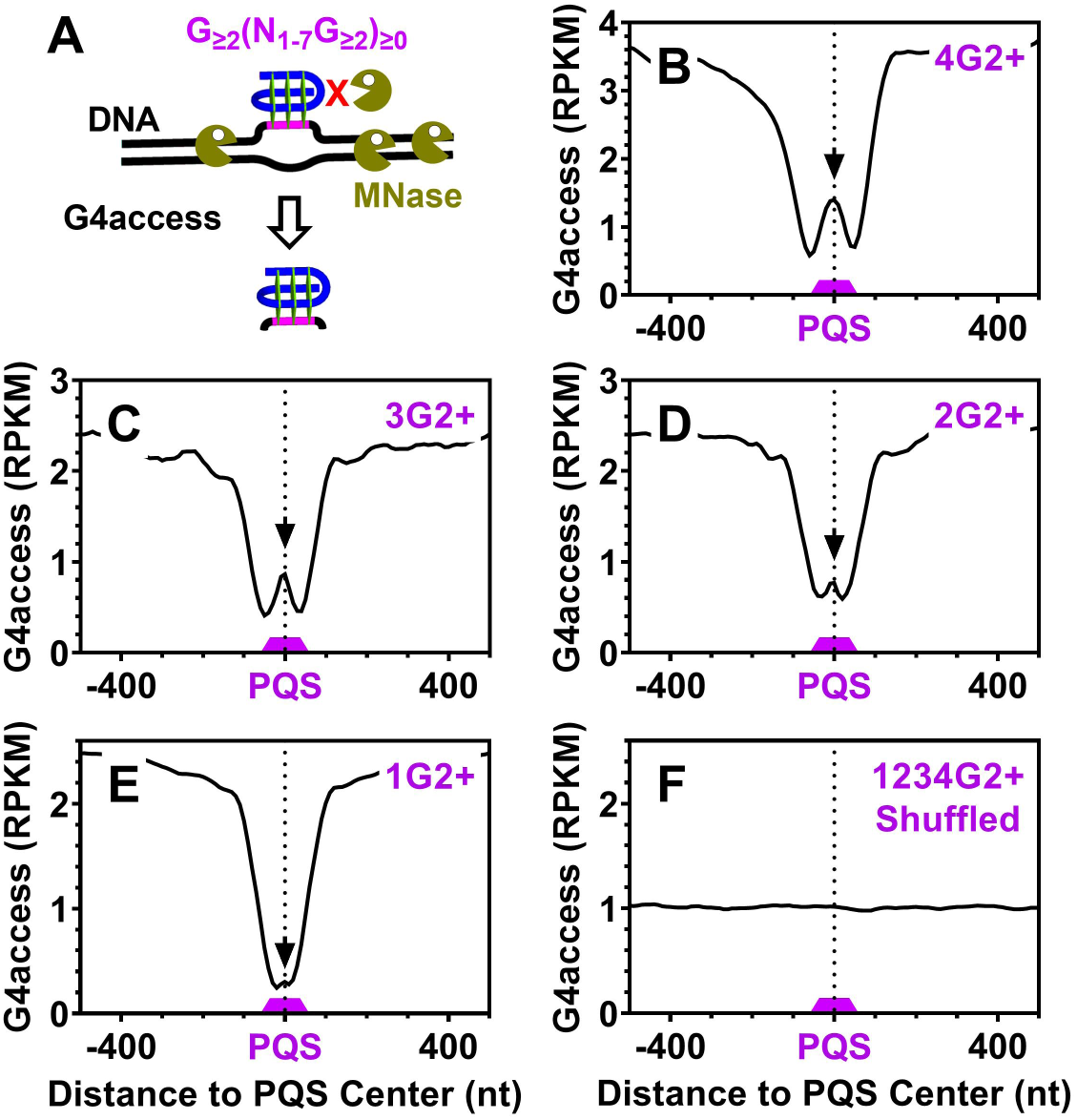
Detection of hG4 formation in the mouse genome by G4access at orphan PQSs containing 1 to 4 or more G_≥2_ tracts that can form G4s of two or more G-tetrads. (A) Schematic illustration of G4 detection using G4access. (B-E) Enrichment of G4access signal at PQSs capable of forming (B) dG4s or (C-E) hG4s. (F) Distribution of G4access signal at randomly shuffled PQSs. The PQSs were flanked on both sides by at least 50 nt of G_≥2_-free regions. Bin size: 10 nt.

